# Using Machine Learning To Develop A Fully Automated Soybean Nodule Acquisition Pipeline (SNAP)

**DOI:** 10.1101/2020.10.12.336156

**Authors:** Talukder Zaki Jubery, Clayton N. Carley, Arti Singh, Soumik Sarkar, Baskar Ganapathysubramanian, Asheesh K. Singh

## Abstract

Nodules form on plant roots through the symbiotic relationship between soybean (*Glycine max* L. Merr.) roots and bacteria (*Bradyrhizobium japonicum*), and are an important structure where atmospheric nitrogen (N_2_) is fixed into bio-available ammonia (NH_3_) for plant growth and developmental. Nodule quantification on soybean roots is a laborious and tedious task; therefore, assessment is done on a less informative qualitative scale. We report the Soybean Nodule Acquisition Pipeline (SNAP) for nodule quantification that combines RetinaNet and UNet deep learning architectures for object (i.e., nodule) detection and segmentation. SNAP was built using data from 691 unique roots from diverse soybean genotypes, vegetative growth stages, and field locations; and has a prediction accuracy of 99%. SNAP reduces the human labor and inconsistencies of counting nodules, while acquiring quantifiable traits related to nodule growth, location and distribution on roots. The ability of SNAP to phenotype nodules on soybean roots at a higher throughput enables researchers to assess the genetic and environmental factors, and their interactions on nodulation from an early development stage. The application of SNAP in research and breeding pipelines may lead to more nitrogen use efficient soybean and other legume species cultivars, as well as enhanced insight into the plant-*Bradyrhizobium* relationship.

## 1. Introduction

The dynamic and symbiotic relationship between soybean (*Glycine max* L. Merr.) and bacteria (*Bradyrhizobium japonicum*) is largely considered mutually beneficial [1]. In a specialized root structure, known as a nodule, bacteria fix atmospheric nitrogen (N2) into a bio-available ammonia (NH3) form that is used by the host soybean plant to assist in meeting its nitrogen needs. The bacteria, in turn, acquires carbon from the host [2, 3]. Nitrogen (N) is critical for building amino acids, vegetative growth, and for protein accumulation in seed. The number of nodules formed on soybean roots can vary from only a few nodules to several hundred per plant [4]. Figure 1 shows diverse soybean genotypes at the Soybean vegetative growth stage (V5) with varying amounts of nodules, which can vary in quantity between genotypes and fluctuate along the taproot and secondary roots of the plant. Comparisons of nodulating versus non-nodulating soybean isolines show there can be a six-fold increase of nitrogen in the nodulating plant at later growth stages, demonstrating the impact of nodulation [5]. However, hyper-nodulating mutants have been shown to be inefficient, with reduced plant health and associated reduced biomass and yield [6].

**Figure 1:**
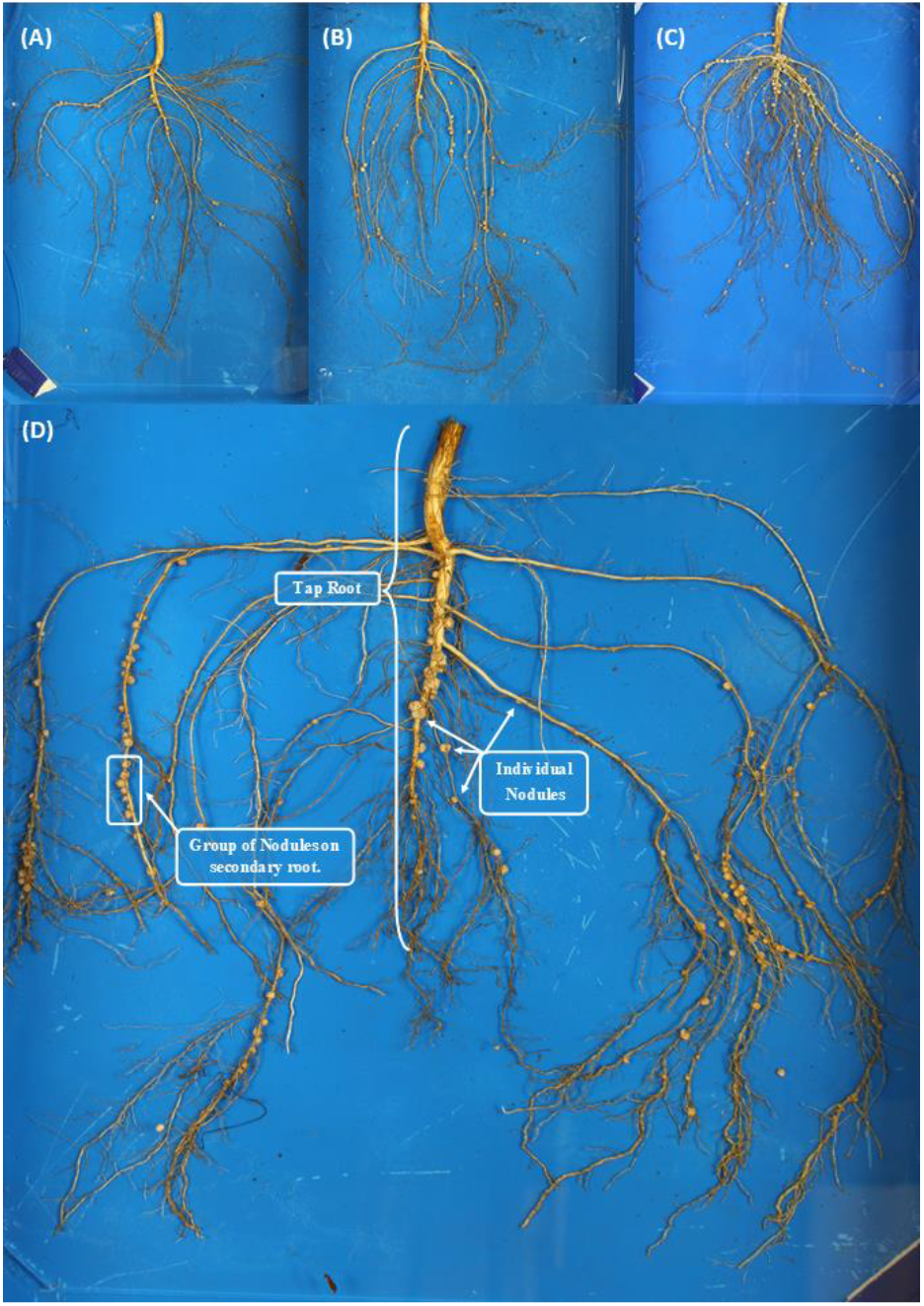
Various genotypes grown in the same field environment with (A) low, (B) moderate, and (C) high nodulation. (D) shows a typical soybean root, with the tap root, scattered (individual) nodules and groups of nodules on plant root.

Due to the importance of nodulation in legume crops in terms of crop health and yield, numerous studies have investigated nodule distributions on roots [7, 8, 9, 10, 11]. To evaluate these nodule plant relationships, nodule evaluations have traditionally been done with qualitative ratings taken on the entire root, just the tap root, secondary roots, or combinations thereof. For the primary world row crops, seed yield is one of the major breeding objectives. However, for N-fixing legume crops, much work is still required to develop and exploit an optimum balance between host genotype, nodulation amount, N rates, and positioning of nodules on the root [12, 13, 14, 15]. Therefore, plant breeders and researchers are motivated to continue the exploration of germplasm, and interactions between the plant-bacteria at multiple levels (cellular, plant, crop, eco-system) [16, 17, 18]. The need to understand nodule growth and development has led to numerous nodulation studies focused on the rates and positioning of biological products [19], fertilizer application patterns [20], climate [21] soil types [22] and even herbicides [23], while evaluating N management decisions including runoff and conservation from previous years production [24] to help mitigate environmental damage and hypoxia zones [25]. Plant breeders can mitigate some of these challenges by developing more efficient and environment responsive N-fixing varieties, to positively impact plant growth and development. For both researchers and producers, a technological limitation is the inability to count and quantify the amount, shape, and location of nodules, as the arduous phenotyping task is very time consuming and can be technically challenging.

Due to the labor-intensive nature of studies on nodule count and its health, researchers have traditionally used qualitative ratings that are based on tap root nodulation representing the entire root, including secondary root branches [26]. Later, more descriptive scales were developed to assist with nodule counting and ratings. These include numerical qualitative scales (ratings 1 to 10), where rating ‘1’ represents few nodules on the taproot and rating ‘10’ with much of the taproot nodulated [27]. In the next rating iteration, nodule quantification included all roots (tap and lateral). There have been attempts to use both nodule count and size and more simple numerical scales to make the rating system more informative [27, 28]. However, the lack of automation has hindered large experiments that use high throughput phenotyping to assess a more exhaustive number of genotypes and often forces much of the distribution of nodule counts into qualitative categories limiting more in-depth statistical analysis and evaluation of diversity that could be possible with quantitative evaluations.

Limited attempts have been made to quantify nodule numbers and size in semi-controlled and especially field environments due to the sheer volume of work and labor required to accomplish root nodule phenotyping at a reasonable scale and time. This severely limits the number of experimental units that are examined. There has been some effort to identify nodulation patterns in legume roots grown in controlled or semi-controlled environments by using traditional computer vision techniques [14, 29, 30]. These techniques involve simple thresholding based segmentation of root from the background using either color alone [14] or color and texture together [29], and then detection of nodules using predefined morphological and color characteristics [14] or predefined outline/shape of the nodules [29]. These techniques were not robust enough to detect all nodules on an image, and users’ input is required where automatic detection fails. Although semi-autonomous counting methods have been developed, full automation for phenotyping is still unavailable, necessitating a reliance on qualitative ratings.

With the advances in phenotyping methods in plant organs [31, 32, 33] and plant stress traits [34, 35, 36, 37, 38], machine learning methods are an attractive solution to advance nodule phenotyping. Machine learning (ML) has been used in numerous plant trait phenotyping to make trait acquisition more feasible and consistent [39]. For example, in disease identification [24, 40, 41, 42], abiotic stress [43, 37], small object detection for SCN eggs [44], and yield-related traits [45, 46]. Furthermore, ML methods have helped develop an end to end phenotyping pipeline to study root system architecture (RSA) traits in soybean [47, 48].

Due to the success of ML for trait phenotyping, we explored ML methods to develop an automated pipeline to identify and characterize nodules and root nodule distributions while reducing the amount of manual processing and counting. The objectives of this work are: (1) to develop an open-source analysis pipeline that can automatically detect nodules on diverse genotypes, and growth stages in soybean root images, and (2) provide statistics including the total number of nodules, sizes of the nodules, and nodules distribution along the tap root. We present a novel Soybean Nodule Acquisition Pipeline (SNAP) to achieve these objectives.

## 2. Materials and Methods

### 2.1 Plant Materials, Root Excavation and Imaging Protocols

The data set for developing SNAP consisted of growing unique soybean genotypes in diverse environments with data collected across several time points. For the evaluation of SNAP, 691 images were collected from 7 unique genotypes (CL0J095-4-6, PI 80831, PI 437462A, PI 438103, PI 438133B, PI 471899, IA3023), in three environments. These included Muscatine Island Research Station, Fruitland IA, in 2018 and 2019 (Soil type: Fruitland Coarse Sand), and The Horticulture Research Station, Gilbert, IA, in 2018 (Soil type: Clarion Loam modified in 1967 to a texture of sandy loam).

Images were collected at three vegetative growth stages: V1, V3, and V5 [49]. Three seeds per experimental unit were planted, and after emergence, two were cut at the soil surface using a sharp blade to leave one standing plant per plot, therefore, each experimental unit consisted of one plant. Each plot was spaced 100 × 100 cm. At the designated growth stage, plants were tagged with barcodes and also labeled with identification strips. Soybean roots were extracted using trenching spades from a 50-cm diameter and 30-cm deep area. Extreme precaution was taken in digging the soil sample to avoid disruption in the plant roots. This was followed by gently removing the loose roots from the soil by hand, ensuring maximum nodule retention on the roots.

After extraction, the root from each plot was placed in a 5-gallon bucket half full of water to rinse the remaining soil from the sample. After 30 minutes, each root was placed on blue painted trays for background consistency. The tray measurements were 35 × 50 cm with a 2 cm lip. To obtain a clear 2D image of each plant root, the imaging trays were half-filled with water, and the roots were gently separated from each other to prevent increased occlusion or clumping together of the roots in each image. Each placement of the root typically took 2-3 minutes. A glass plate fitted to the size of the tray was then laid on top of the root in the water to hold it in place and then slid into an imaging platform [48] customized for this project. The platform was built from aluminum T-slot extrusion frames (80/20 Inc., Columbia City, IN) with two softbox photography lights (Neewer; Shenzen, China), four 70-watt CFL bulbs in total, to provide consistent illumination and a Canon T5i digital SLR camera (lens: EF-S 18-55mm f/3.5-5.6 IS II) (Canon USA, Inc., Melville, NY) was mounted 55cm above the imaging plane. See Falk et al. [48] for full details. The camera was tethered to a laptop with Smart Shooter 3 photo capture and camera control software [50], to trigger image capture. Smart Shooter enabled automatic image labeling and naming by reading the tag barcode in each image, reducing human error.

After imaging, roots were dried in paper bags at 60°C for two days. After the roots were thoroughly dry, they were weighed for dry weight (grams), and nodules from each root were manually removed (by hand) and counted for use in the ground truth analysis. Hand removal of the nodules was accomplished by trained researchers, who carefully removed each nodule with tweezers. Upon removal, a second researcher cross-validated and observed the root for any remaining nodules. This two-person unit then individually counted and recorded the number of nodules. If there was a discrepancy in the counts, that sample was recounted, ensuring that all nodules were correctly identified and counted. The removed nodules were then weighed and recorded in grams. After completing a sample and its validation, the two-person team moved on to the next sample.

### 2.2 Deep Learning and Image Processing Workflow

The proposed workflow using deep learning (DL) and image processing is shown in Figure 2. It has two phases: 1) training, and 2) evaluation. Two DL networks were trained: (a) nodule detection network, and (b) tap root detection network. To train these networks, we selected a representative subset of the data set for annotation using submodular optimization [51]. Nodules were annotated by a human root nodule phenotyping expert, who drew bounding boxes around each nodule on 256 × 256 image patches using VGG annotator [52] (Figures S.1–3). The taproot was annotated using the full root image, and tracing was done on a Microsoft Surface computer with a Surface pen generating a line over the tap root using the edit and create tool in Microsoft Photos (Microsoft 2020). In the evaluation phase, we used the trained models to obtain the number of nodules, size distribution, and spatial distribution along the tap root.

**Figure 2:**
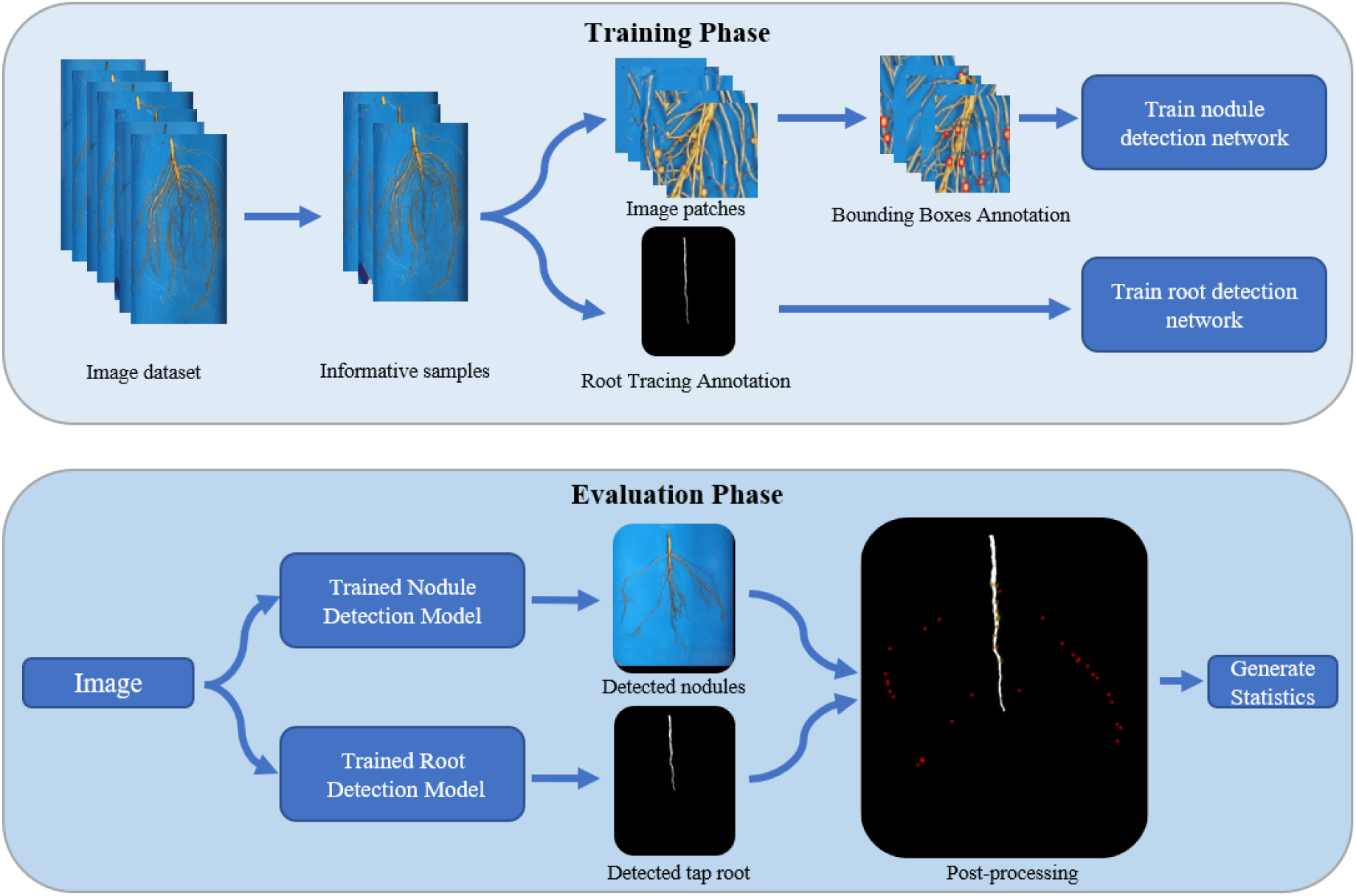
Overview of the workflow of Soybean Nodulation Acquisition Pipeline (SNAP). The top box contains workflow for the training phase, including training of nodule detection and taproot detection. The bottom box contains workflow for the evaluation phase, including the evaluation of nodule locations and metrics and in relation to detected tap roots.

#### 2.2.1 Representative Sample Selection

We deployed an unsupervised framework based on submodular function optimization [53] to select a representative subset (R) from the whole data set (W) for annotation. One of the properties of this function is that incremental gain of utility by adding new samples to the subset (R) decreases as the size of the subset grows. Thus, a small subset is maximally representative of the full dataset. We used the uncapacitated facility location function [54] as the submodular function:

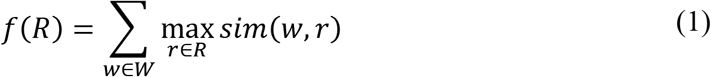

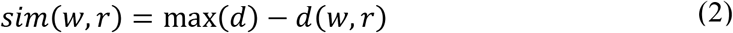

Where *sim*(*w,r*) and *d*(*w,r*) are a measure of similarity and Euclidean distance between a pair of samples *w* ∈ *W,r* ∈ *R*. The value of *f*(*R*) indicates the representativeness of the subset *R* to the whole set *W*. By maximizing *f*(*R*), we selected the most representative subset *R*, given that |*R*| = *K*, where *K* is a predefined value indicating the size of subset. We used a simple greedy forward-selection algorithm [55] to maximize *f*(*R*). The algorithm starts with *R* = ∅ and iteratively adds samples *r* ∈ *W* \ *R* that maximally increases *f*(*R*).

In the framework, each sample with dimensions 5000 × 3500 × 3 is input as a vector. In high dimensional space, Euclidean distance becomes less meaningful [56]. Therefore, we reduced the dimensions of the input vector using a combination of nonlinear transformation (an autoencoder) to balance an accurate reconstruction without overfitting the training data and linear transformation (principal component analysis (PCA)) to best minimize feature redundancy. First, we down-sampled the input image (5000 × 3500 × 3) using bi-linear interpolation (to 512 × 512 × 3), ensuring no visible distortion of the root. Next, we generated low-dimensional latent representation using a shallow autoencoder, then, we flattened the encoded representation and finally mapped it to 400 principal components that capture approximately 98% of the variance (Figure S.4).

#### 2.2.2 Nodule Detection Framework

We approached nodule detection as a dense detection problem, as the root images have many target nodules for detection. We selected RetinaNet [56], which shows better performance than other detection frameworks for dense detection problems due to the use of “Focal Loss” as a classification loss function [56]. Focal loss function uses a scaling factor to focus on the sparse set of hard examples (foreground/nodules) and down-weights the contribution of easy (and abundant) examples (background) during training (Figure S.5).

We used the same backbone and subnetworks as Lin [56]. The backbone network was based on ResNet50-FPN and two subnetworks consisting of four 3×3 convolution layers with 256 filters, Rectified Linear Unit (ReLU), and sigmoid activation functions. Multiple anchors were used at each spatial location to detect nodules of various sizes and aspect ratios. The default anchor configurations in Lin [56] were suitable for detecting objects with 32 pixels or above in size. We changed the default configurations and optimized them for our case. We explored two different selection strategies: a) maximize overlapping between selected anchors and bounding boxes as Zlocha [57] using a differential evolution search algorithm [58], b) fit a normal distribution to the bounding box sizes and aspect ratios and make selections based on equidistant percentiles. In our case, we evaluated equidistant percentiles using several scales and aspect ratios of 3, 5, and 7, where the default anchor configurations consisted of three scales (2^0/3^,2^1/3^, and 2^2/3^) and three aspects (1:2, 1:1, and 2:1).

The base ResNet-50 models were initialized as the pre-trained model in the Coco data set [59]. The convolution layers on the subnets, except the last one, were initialized with bias b = 0 and a Gaussian weight fill with *σ* = 0.001. The last layer on the classification subnet is initialized with *b* = −*log*(1−*π*)*/π*. We set *π* = 0.01, where *π* specifies the confidence value of an anchor as foreground.

Data augmentation was performed by first flipping the images in horizontal and vertical directions with a 50% chance. Next, random affine transformations with rotation/shearing of up to 0.1 radians were used, followed by scaling/translation of up to 10% of the image size. To develop the trained networks and evaluate them, we utilize a standard 80% training and 20% testing set. The selections were made before data augmentation.

The model was trained with an Adaptive Moment Estimation (Adam) optimizer [60] with an initial learning rate of 10^−3^. Focal loss was used to train the classification subnetwork.

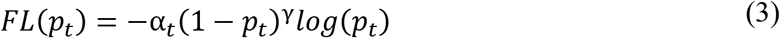

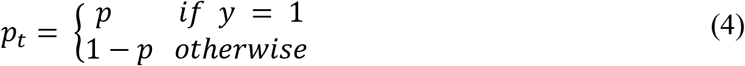

Where *α*_*t*_ and *γ* are scaling factors to focus on the sparse set of hard examples (nodules). *p* is the class probability for the hard examples, *y* = 1. We used *α* = 0.25 and *γ* = 2.0 [56].

The standard smooth L1 loss was used for box regression.

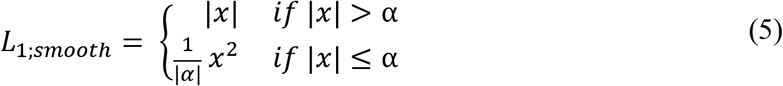

With *α* = 1/9. *α* is the transition point (near zero) from L1 to L2 loss.

The number of trainable parameters of the framework was 36,737,717. We performed data augmentation and explored the effect of batch size and scale of input image size on our dataset. The average precision (AP) [61] metric was used to evaluate the models. The AP indicates the area under the precision-recall curve after samples are ordered based on the confidence of detection.

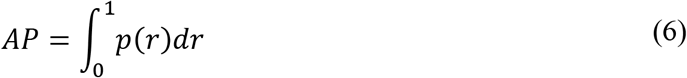

Where p(r)is the precision as a function of recall.

All models were trained for 300 iterations on fixed training data (representative 20% samples from the dataset) and tested on a fixed test-data using 4% randomly selected samples from the dataset. Model development in the pipeline was completed using a GeForce GTX TITAN X 12 GB GPU. On average, the training time took from 10 hours to 3.5 days for the models.

#### 2.2.3 Tap Root Detection Framework

The tap root detection was approached as an image segmentation task. We deployed a Unet involving three 2 × 2 maxpooling operations (down sampling) [62] with 7,708,609 trainable parameters. The network consisted of three encoding/contracting blocks and three decoding/expansive blocks. Each encoding block consisted of two 3 × 3 convolutions with 64 feature channels, two rectified linear unit (ReLU) activations, two batch normalizations, and one dropout operation. The decoding block was the same as the encoding block, except for the dropout operation. In between these encoding and decoding blocks, the output from the encoding block was down sampled using a 2 × 2 maxpooling operation followed by two 3 × 3 convolutions with 128 feature channels, two rectified linear unit (ReLU) activations, two batch normalizations, and two dropout operation. The output from these operations was up-sampled using a 2 × 2 transposed convolution, followed by a concatenation operation that combines the encoder output feature channels and up-sampled feature channels. After the decoding blocks, the final layer is a 1 × 1 convolution that is used to map each of the 64 channel feature vectors to the 1 channel output binary image (Figure S.6).

We utilized a standard 80% training and 20% testing set. The selections were made before data augmentation. Data augmentation is performed by flipping the images in horizontal and vertical directions, zooming 120%, translation in horizontal and vertical directions then 5%, rotation until 15 degrees.

The model was initialized using Glorot uniform initializer [63] with zero biases and trained with Adam optimizer [60] with batch size 4 and learning rate 10^−3^ using a Jaccard index [64] as the loss function (Figure S.7).

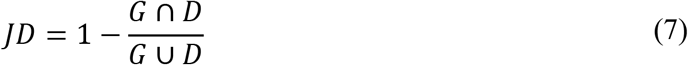

Where *G* is the annotated ground truth tap root image, and *D* is the detected tap root image.

### 2.3 Hyper-parameters tuning

Of the 20% representative samples used for training-data, 4% were randomly selected as test data. During hyper-parameter tuning, model training was performed on the represented samples, and model evaluation was done on the test data.

### 2.4 Detection, Post-Processing and Evaluation

In the trained nodule detection model, each sample was fed to the trained RetinaNet model in 256 by 256 patches. Samples were padded to ensure the image width and height were divisible by 256.

In order to begin elucidating spatial relationships of nodules in various root zones, we developed a method of predicting the number of nodules in the taproot location zone using a trained Unet as shown in Figure 3. Once the model identified the taproot location in the image, we dilated the taproot. We then identified the center of the detected bounding boxes around nodules which fell within the dilated taproot. We then count the bounding box centers that are within the dilated taproot zone and called these as taproot zone affiliated nodules. Further, in order to generate statistics involving the spatial distribution of the nodules along the taproot, we skeletonized the taproot and identified the nearest location on the taproot for every identified bounding box. This enabled spatial statistics assessments of nodules along the taproot related in proximity to the soil line and taproot length.

**Figure 3:**
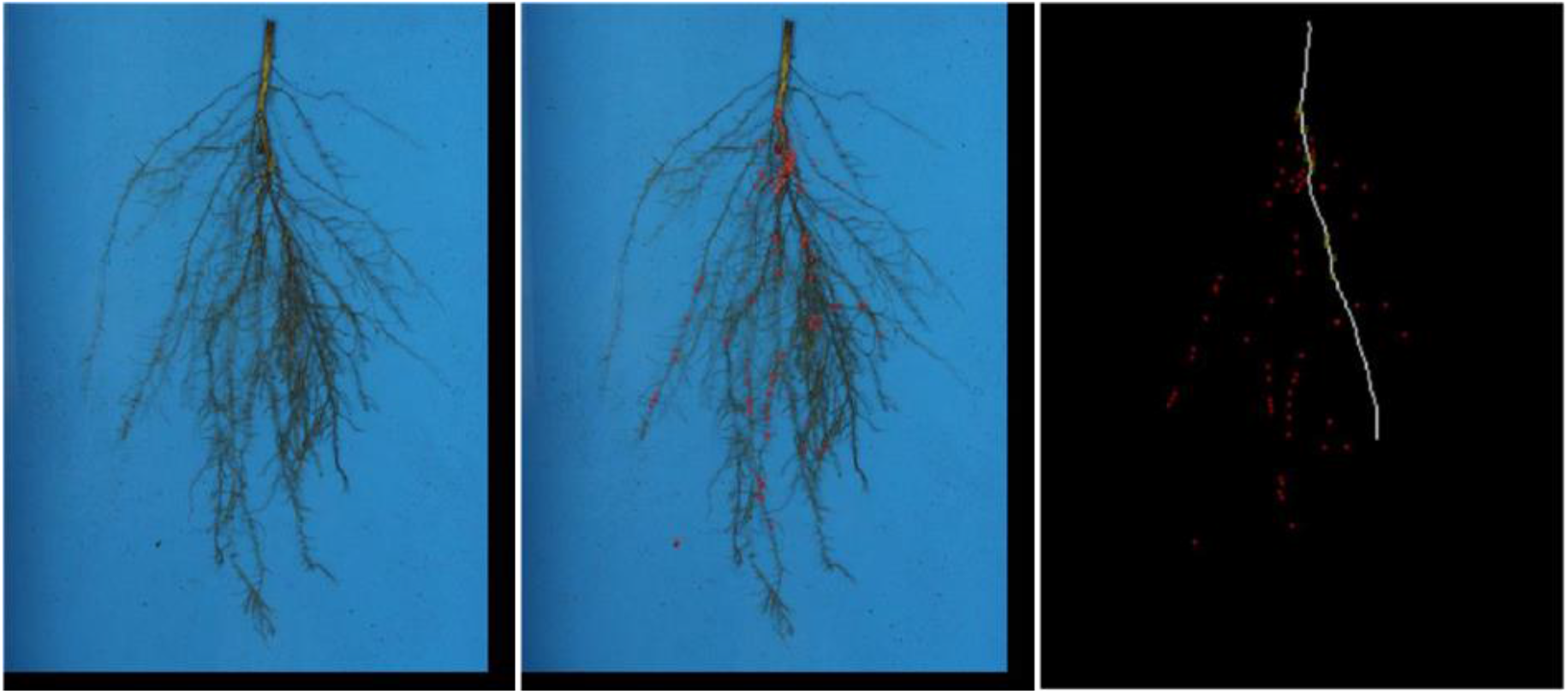
Left) Sample input image; Center) SNAP output image with bounding boxes around predicted nodules; Right) SNAP output image with detected taproot (white) and locations of the center of the bounding boxes (red points). Nodules with the center of its bounding box within the taproot region were considered adjacent or connected to the taproot (green points).

### 2.5 Avoiding Misclassification of Nodules and SCN Cysts

To avoid potential misclassification of early developing nodules with additional structures such as Soybean Cyst Nematode (SCN) (*Heterodera glycines*) cysts, during the training phase, images with SCN cysts on the roots were included in the training dataset to enable robust, accurate classification of only nodules. When labeling nodules, care was taken not to mislabel a cyst as a nodule. To evaluate the accuracy of SNAP predicted nodules, a human expert rater evaluated each predicted nodule to ensure that it was not a cyst.

### 2.6 SNAP Evaluation

Further evaluation of SNAP was conducted to evaluate the sensitivity and precision of the pipeline nodule predictions using the following equations:

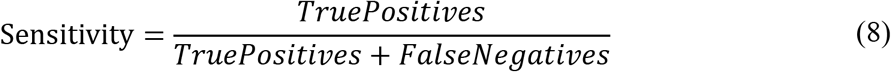

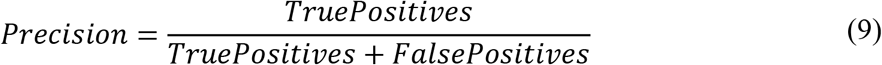

Where true positive represents instances when SNAP accurately identified nodules in the image, false negative when nodules were present in the image but not identified by SNAP, and false positives were twice counted nodules or the predicted nodule was not an actual nodule.

To evaluate the required processing time of SNAP, the pipeline was implemented using a Python 3.6 environment on a Microsoft surface with 16 GB ram using an Intel(R) Core(TM) i7-8650U CPU @ 1.90GHz 2.11GHz.

## 3. Results

To develop a model that best quantifies nodules, a balance between computational resources and accuracy was sought. When evaluating accuracy, an increase of 20% average precision (AP; equation 6) was observed for the optimized method compared to the default, and minimal AP difference was noted when the number of scales and aspect ratios were increased (Supplemental Table S.1). The normal percentile method was computationally cheap, compared to the optimized method, and it enforces the expected normal distribution on the naturally occurring objects like nodules. No further improvement was noted for the normal percentile method with increasing scales and aspect ratios. We investigated the effect of % data annotated, batch size, and input image scale by comparing AP (Table 1).

**Table 1:**
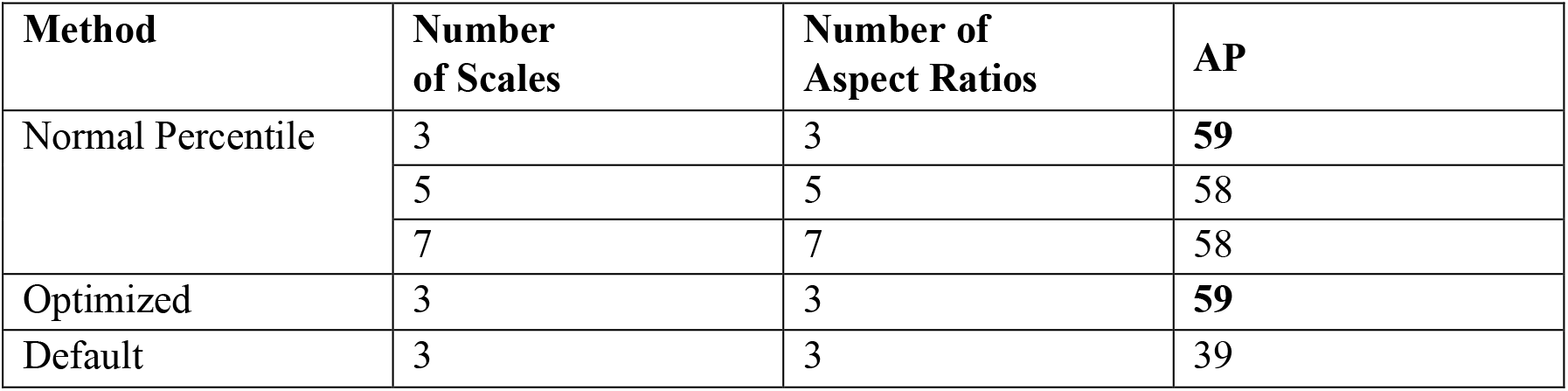
Effect of anchor selection method sizes and aspect ratios on Average Precision (AP) for SNAP evaluation of individual ML nodule detection models.

No perceivable difference was noted on the effect of batch size. Minimal improvement was noted with the increase of percent data annotated, except at 30%, a small increase in AP was noted with an increase in input image scale. (Table 2). No major improvement was noted when the batch size was increased to 32 and 64, and the input image scale was increased to 758. The image input scales tended to over-predict nodules bigger than about 15 pixels, which roughly represents the width of the bounding box (Figure S.8). An improvement in nodule detection was noted for smaller sized nodules (<15 pixels), when the input image scale was increased from 256 to 512 scale without a continual improvement from 512 to 768 scales.

**Table 2:** Effect of batch size and input image scale for soybean nodule detection at varying percentages of annotation.

| % Data (Annotated) | Batch Size | Input Image Scale | AP |
| --- | --- | --- | --- |
| 10 | 8 | 256 | 55 |
|  | 16 | 256 | 56 |
|  | 8 | 512 | 58 |
|  | 16 | 512 | 58 |
| 20 | 8 | 256 | 57 |
|  | 16 | 256 | 59 |
|  | 8 | 512 | 59 |
|  | 16 | 512 | 59.5 |
| 30 | 8 | 256 | 57 |
|  | 16 | 256 | 59 |
|  | 8 | 512 | 60.5 |
|  | 16 | 512 | **62** |

### 3.1 Validation of whole root and tap root nodule counts

To determine the capability of SNAP for nodule detection, we randomly picked 10% of the root sample images not used in the ML model training and evaluation sets. The validation was performed three ways by comparisons between; (a) SNAP nodule count to human rater removed nodule count, (b) SNAP nodule count to human expert nodule count from the image, and (c) SNAP nodule count in the tap root zone to human nodule count on the taproot zone in the image (Figure 4). Examples of good and poor nodule predictions can be found in Figures S.9 and S.10.

**Figure 4:**
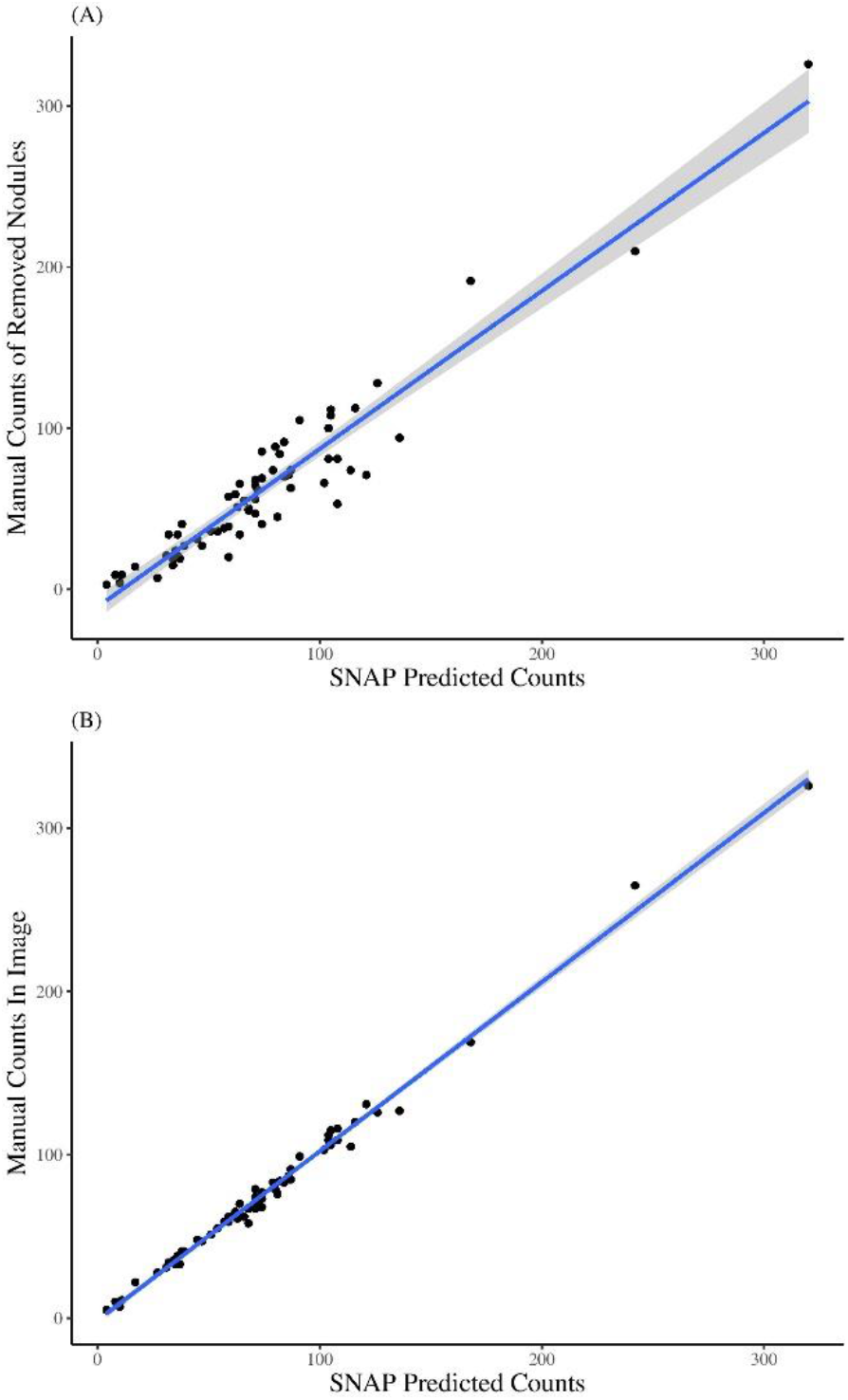
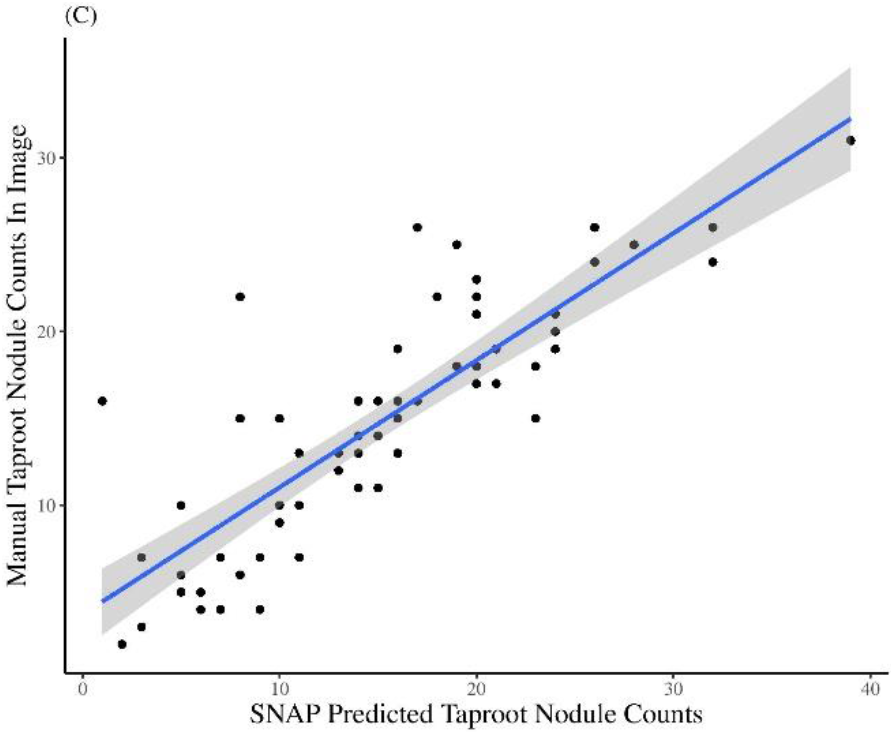
(A) Manually removed and counted nodules vs SNAP counted nodules. (B) Image counted nodules vs SNAP counted nodules. (C) Tap root image counted nodules vs SNAP counted nodules in the tap root zone.

High correlations were observed for all the three comparisons, with R^2^ of 0.91 in the physically removed nodules to SNAP comparison, 0.99 in the nodules counted within the image to SNAP comparison, and 0.71 in the root zone counted nodules within the image to SNAP taproot zone nodule counts. Overall, SNAP nodule count had sensitivity = 0.934 and precision = 0.951.

### 3.2 Time and labor requirements

SNAP pipeline development was dependent on efficient root digging, sample preparation, including root washing, imaging, and generation of ground truth data by manual nodule harvesting and counting. Once the ML model was developed, the actual time to obtain nodule count through SNAP was dramatically reduced. Manual nodule harvesting (i.e., extraction) and sample preparation with imaging time increased per growth stage. The most time-intensive step was manual quantification (i.e., ground truth nodule counting), and the time required to remove nodules dramatically increased per growth stage. Once the ML model is trained, the most time-intensive step of manual nodule count is removed, providing SNAP users with an increase in time and resource efficiency, and an ability to work with more samples. In the course of this study, we observed that on average the manual extraction of roots takes 240, 360, and 420 seconds for V1, V3, and V5 roots, respectively. To wash and image V1, V3, and V5 roots, it took an average of 100, 128, and 150 seconds, respectively. The comparison of hand quantification of nodules and SNAP showed a dramatic change, as it took trained workers an average of 1500, 2100, and 3000 seconds per V1, V3, and V5 root, respectively; while SNAP took 90, 120, and 150 seconds per V1, V3, and V5 root, respectively (Table S.2).

## 4. Discussion

Object detection in cluttered and complicated backgrounds is an inherently challenging problem. The complexity and diversity of roots and nodules combined with root occlusion and color/texture similarities of the roots and nodules and the need for a high throughput method to routinely screen large number of genotypes necessitates a unique ML architecture to extract and quantify nodule counts and sizes. Our earlier iterations to approach this problem included segmentation and detection using classical SURF and SIFT methods [65], and a deep learning based Faster-RCNN approach [66]. However, due to poor model performance with these methods, we transitioned to RetinaNet, which showed improved accuracy and faster performance in dense object detection due to the use of focal loss function [56].

We combined RetinaNet and Unet, to develop SNAP that enables accurate nodule counts (91% of manual removal counted and 99% for image counted) on a soybean root and generate information on the spatial distribution of nodules. SNAP provides automation in phenotyping, with a significant reduction in time to count nodules on a root. In each image, nodules were counted in about 2-3 minutes compared to another existing semi-automated pipeline, which took about 20 minutes to do similar counting [14]. The primary reduction in time was observed in SNAP compared to manual counting, with improvements by factors of 16 to 25 times depending on the growth stage.

SNAP offers multiple avenues for its applications in research and breeding domains. There is an active interest in learning the spatial distribution and temporal development of nodulation in crops, particularly to optimize plant performance and symbiosis with bacteria [67, 68]. SNAP can estimate the number of nodules in the taproot zone with a precision of over 70%. Upon human validation of SNAP predicted nodules, no instance was noted where an SCN cyst was misclassified as a nodule. Figure 5 shows a representative example of a complex root architecture with varying nodule sizes, and nodule and cyst distribution patterns. As SNAP is able to identify even small or newly developing nodules often missed in rater assessments, it is possible to now classify nodule development stages and quantities in correlation to vegetative growth stages or evaluate the effects of SCN on nodulation in a temporal scale using a fully automated ML cyst detection pipeline [44]. However, it is important to note that field root study samples are destructively sampled, therefore the study of nodules will require the evaluation of separate plants of the same genotypes at different time points. Through SNAP, the groundwork has been laid for future studies that can screen large breeding populations, identify and investigate QTL, and determine the relationships and correlations between root growth zones, RSA, and nodules.

**Figure 5:**
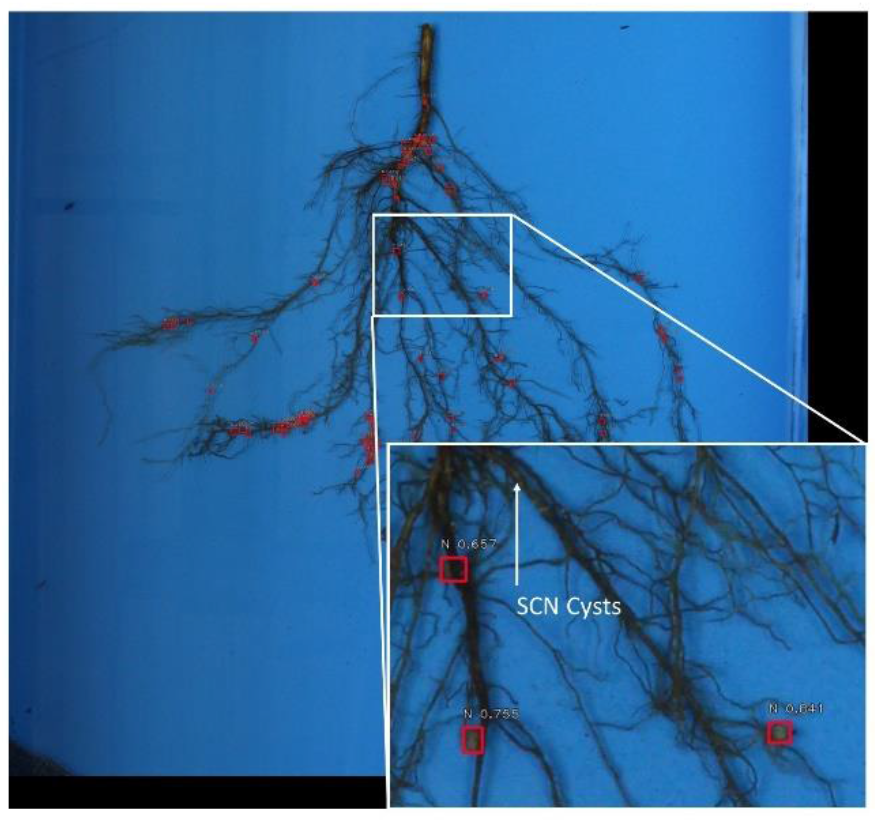
Image of a soybean root sample where soybean cyst nematode (SCN) cysts were present. SNAP did not detect cysts as nodules, showing its robustness.

While we utilized root samples from field experiments, SNAP can be combined with additional technologies such as mobile platforms for immediate in-field evaluation, or non-field environments, such as greenhouse, growth chamber, X-ray, and CT scan experiments to enable further solutions to breeding challenges for nodule phenotyping. SNAP based nodule counting is amenable with previously used methods such as a binned rating scale of 1-10 if researchers are interested in comparative studies combing old and new research outcomes. Using the distribution generated by SNAP, a more accurate binning and count can occur, and roots can be rated automatically for comparison to each other and potentially against additional or prior studies.

Often breeders are unable to include root architecture and nodulation in their assessments as they are seen as unattainable and unrealistic traits to evaluate in a manageable and high throughput manner; although more recently improvements have been suggested [47, 48, 69, 70]. SNAP empowers breeders to evaluate and select genotypes that have a required level of nodulation in various biotic and abiotic conditions and accounting for genotype by environment interactions. Additionally, SNAP increases opportunities to identify and map genes controlling nodule related traits, for example, size, onset, and nodule growth coinciding with plant growth and development stages. Since SNAP was trained and evaluated on several genotypes, field locations, and vegetative growth stages, it can enable the investigation of nodulation across diverse root types and vegetative time points as well as the investigation of the growth of nodules between similar roots in a temporal scale, unraveling new scientific insights at a larger scale (i.e., more genotypes) which was previously difficult.

Non-soybean crop researchers working in other N-fixing crops need to validate the results of SNAP prior to its usage in their research. While we tested the success of SNAP in correctly identifying nodules discriminatively from SCN cysts, there may be other pathogen organisms, for example root knot nematode, that will require additional model training and testing prior to its deployment to study root nodules. With advances in higher resolution imaging, a SNAP type of approach in the future will be beneficial to study other beneficial plant and micro-organism interactions, such as Arbuscular mycorhizal fungi, which can positively impact crop production [71, 72]. The combination of SNAP based nodule phenotyping in conjunction with genomic prediction forecasted on a germplasm collection is also an attractive approach to identify useful accessions for plant breeding applications spanning various maturity groups [73]

Improvements in SNAP functionality could be realized, for example, through the implementation of more sophisticated active learning-based representative sample selection strategy to help improve the performance of the pipeline [74], delineate nodules specifically for irregular and non-uniform non-spherical nodules to get even higher size and shape accuracy, or evaluate spatial distribution of the nodules along the lateral roots.

Overall, SNAP will help reduce the strain on human labor and capacity to quantify nodules consistently and accurately in N-fixing crop species and move the current state of the art in nodule phenotyping and associated applications. SNAP outputs will have usefulness for researchers and farmers, who have an interest to rapidly and accurately phenotype nodules on roots.

## Acknowledgments

The authors thank the many undergraduate, graduate students and staff in the Singh group at Iowa State University who helped with field experiments, data collection and imaging. Additional thanks to Koushik Nagasubramanian for his initial pipeline suggestions, and Vahid Mirnezami and Kevin Falk for assistance with the imaging system.

## Author contributions

C.C., Z.J., B.G., and A.K.S. conceived the project. All authors participated in the project implementation and completion. C.C. conducted experiments, imaging, and data curation. Z.J developed the machine learning and image analysis pipeline. C.C. annotated the ground truth images and assessed the pipeline output. C.C. and Z.J. wrote the manuscript draft with A.K.S and B.G. All authors contributed to the final manuscript production.

## Data Availability

Data is freely available upon request to the corresponding authors, and the pipeline software and codes will be available at GitHub: (https://github.com/SoylabSingh/SNAP).

## Supplementary Materials

**Figure S.1:**
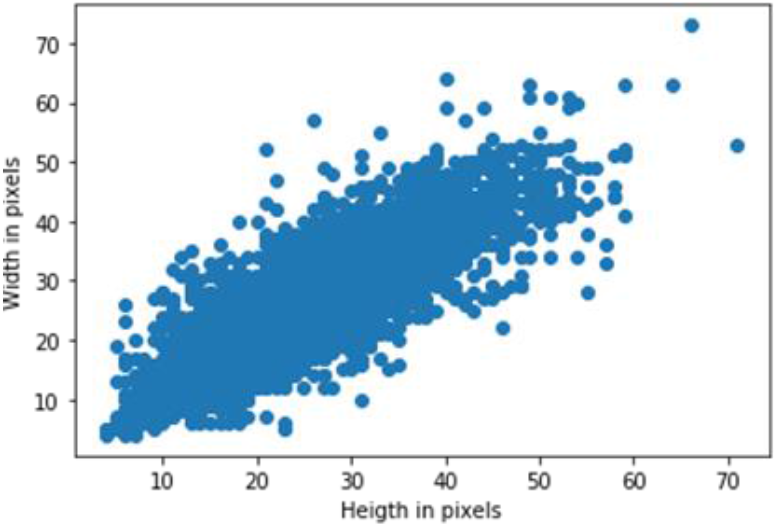
Width and height of the annotated nodules.

**Figure S.2:**
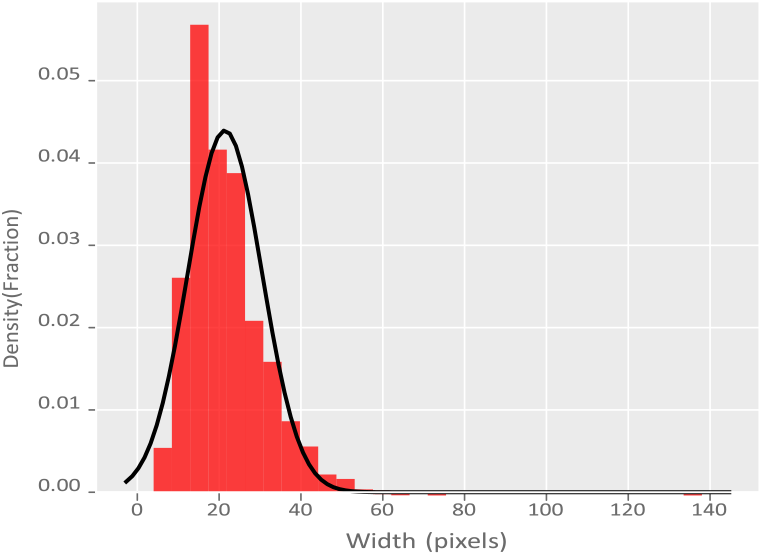
Width distribution of the annotated bounding boxes for nodules from 30% of the dataset.

**Figure S.3:**
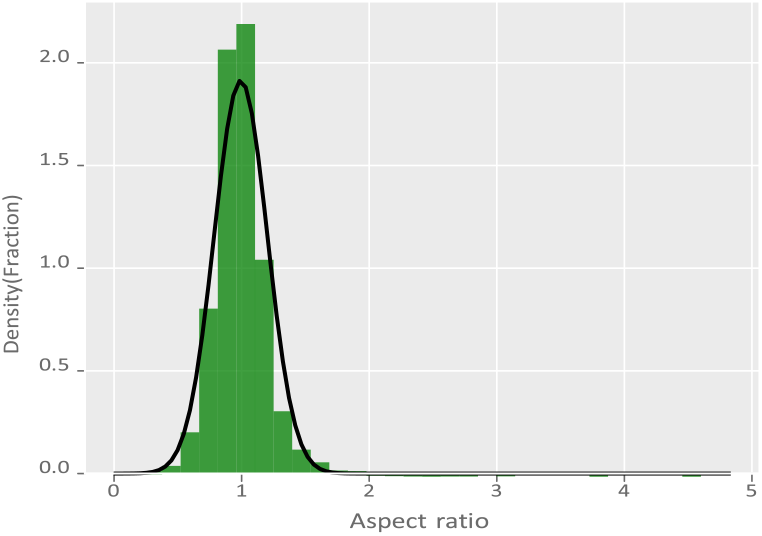
Aspect ratio distribution of the annotated bounding boxes for the nodules from 30% of the dataset.

**Figure S.4:**
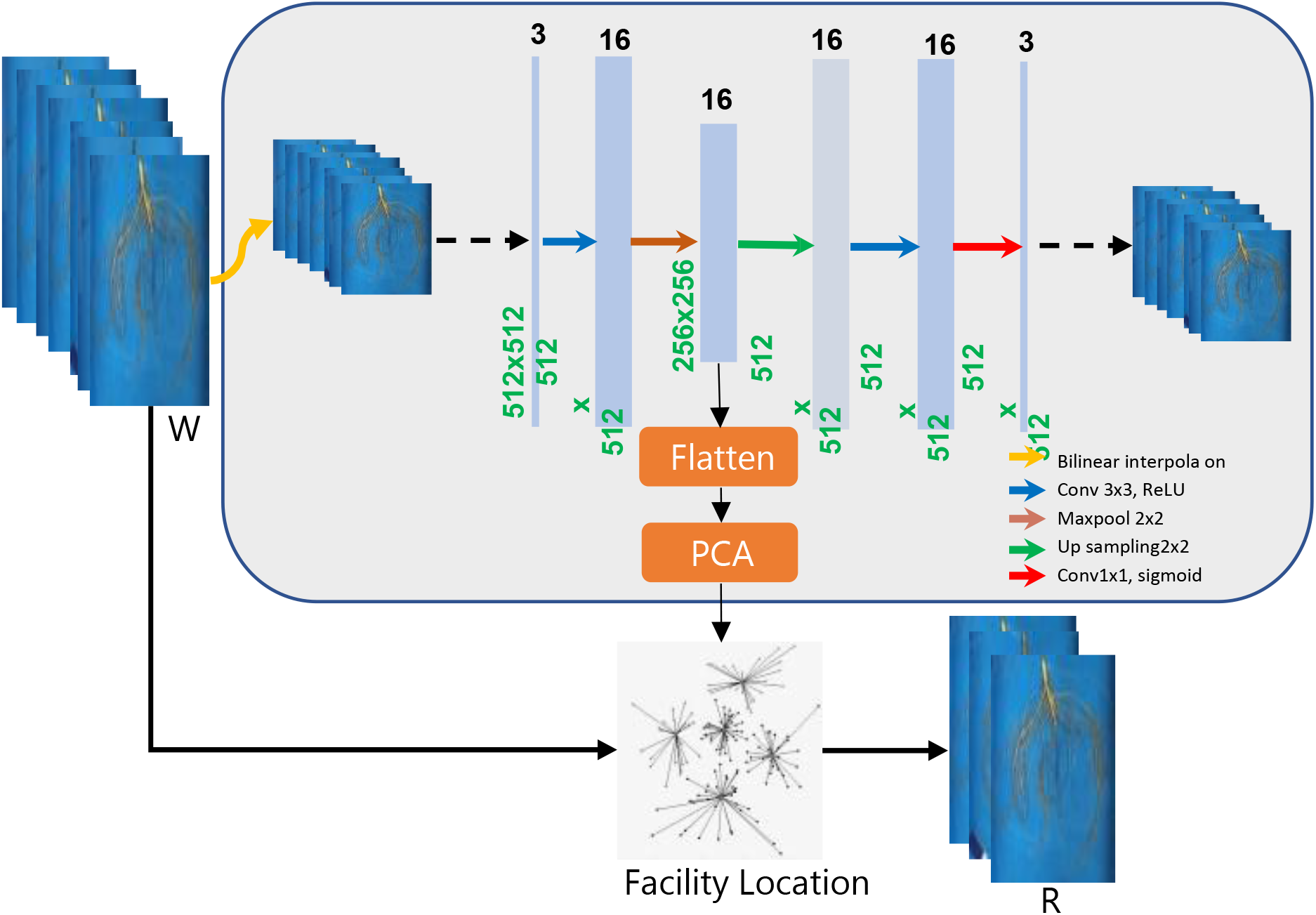
Informative sample selection workflow. Latent features of the images (W) were obtained via dimensionality reduction using convolution autoencoder and principal component analysis (PCA) as shown in the box. A set of information samples (R) (with predefined size) were selected using a facility location based submodular function optimization.

**Figure S.5:**
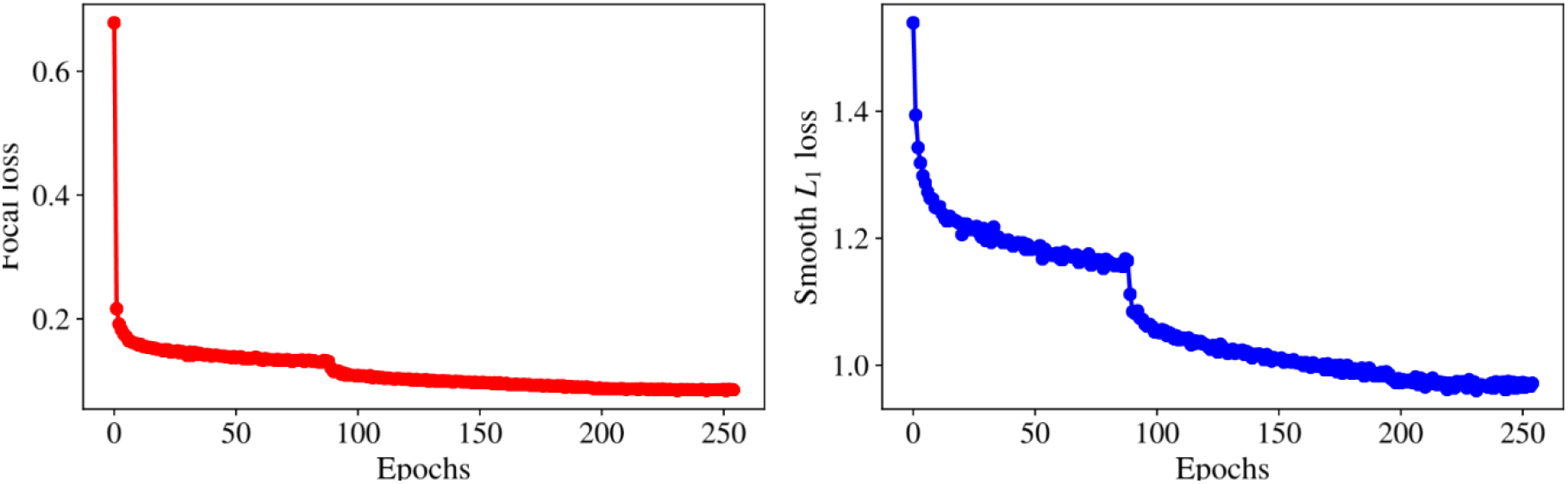
Focal (classification) and regression (bounding box detection) losses during training of nodule detection network, RetinaNet, using input image scale 512, anchor scales 0.48, 0.67, 0.86, aspect ratios 0.85, 0.99, 1.13.

**Figure S.6:**
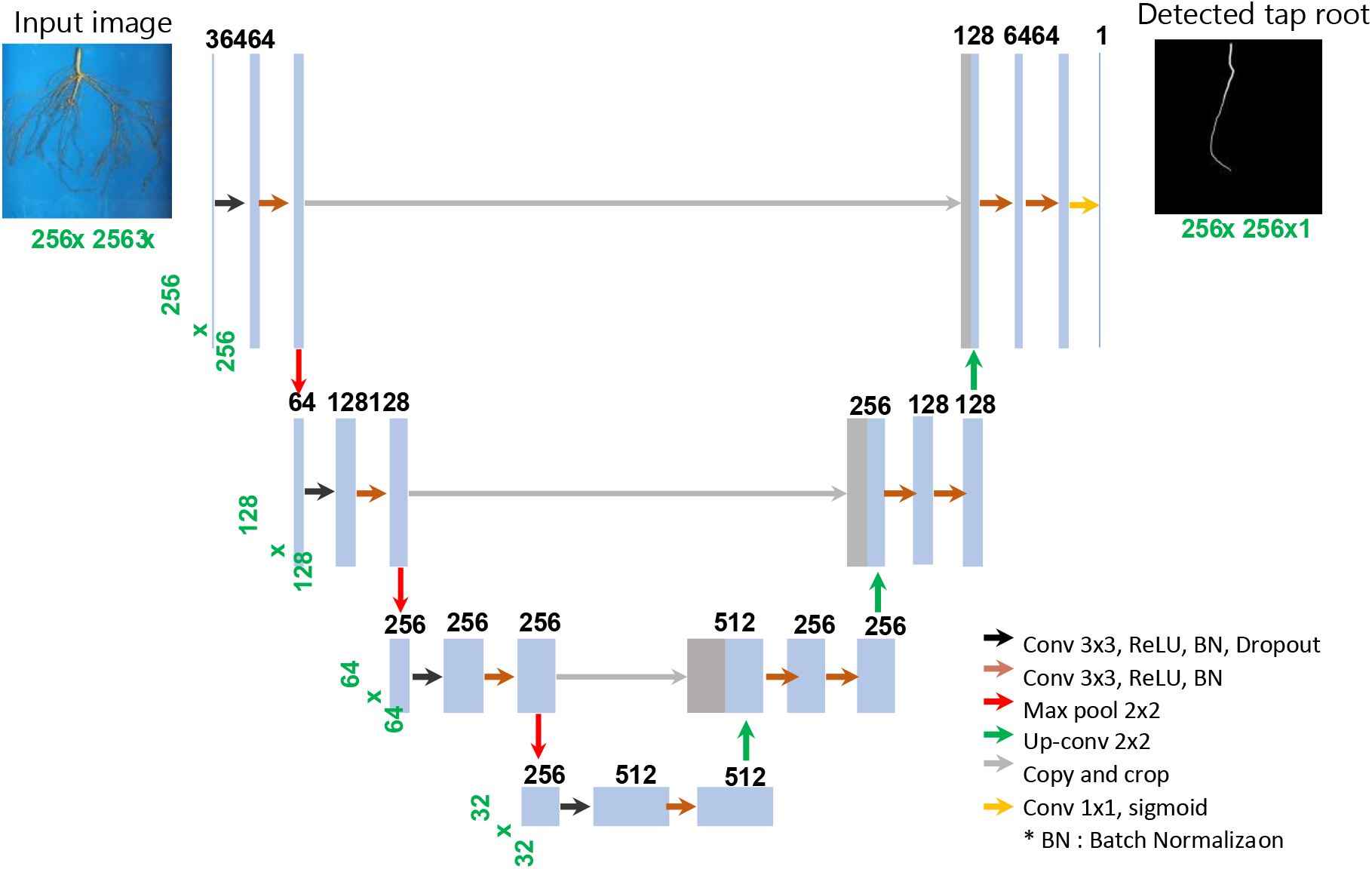
The U-Net architecture used to develop the tap root detection model. Each blue bar represents a multi-channel feature map. The number of channels is denoted on top of the bar, and width × height is provided on the left of the box. Gray bars are copied feature maps. Colored arrows denote the different operations as indicated in the lower right arrow legend.

**Figure S.7:**
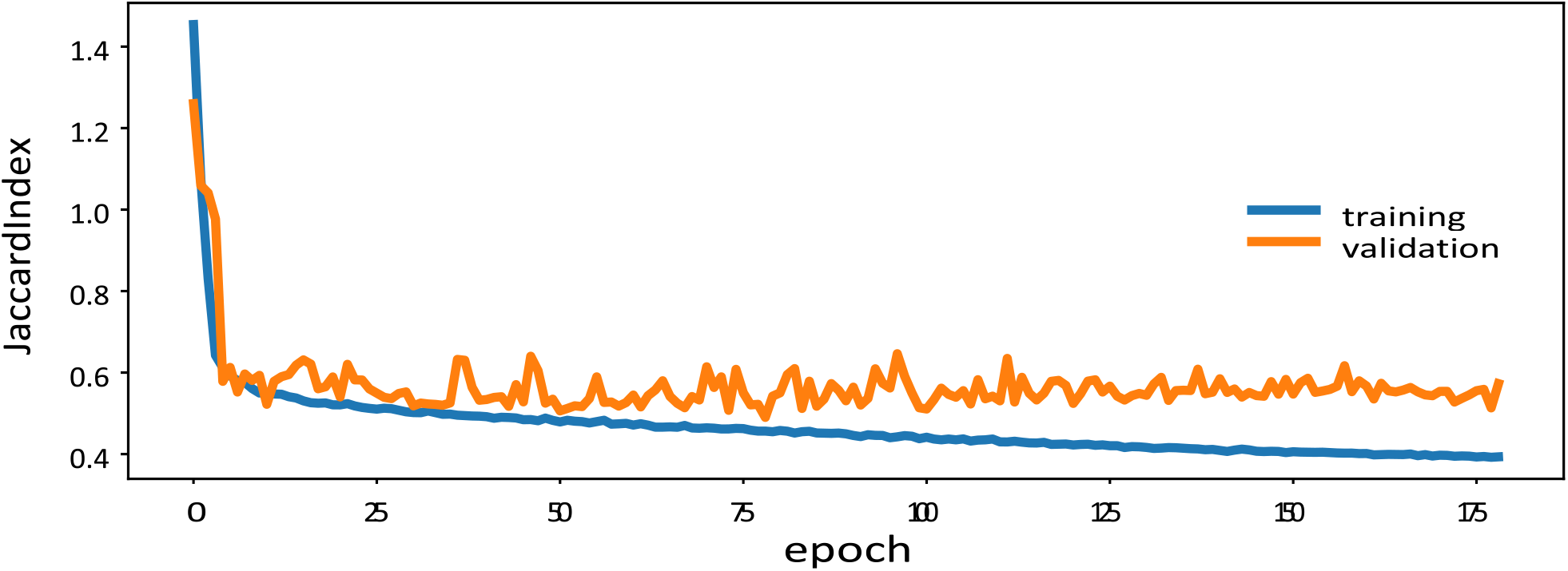
Training and validation losses (Jaccard loss) during the training of U-Net for tap root detection.

**Figure S.8:**
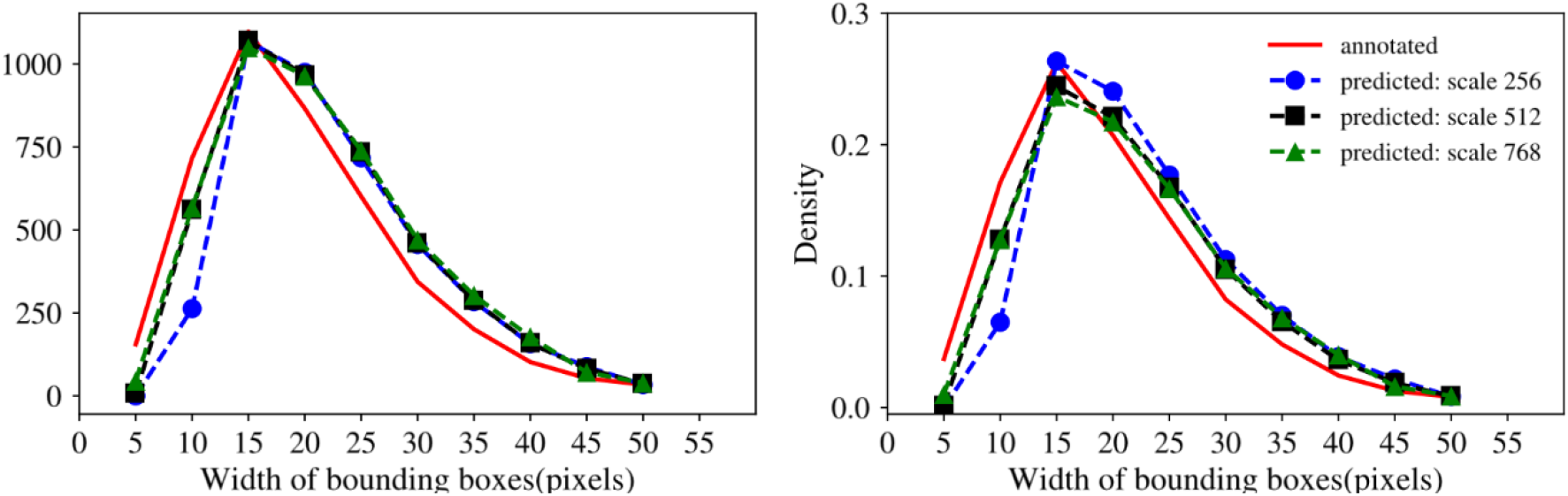
Effect of input image scale on nodule detection in the test data. The red solid line indicates the distribution of the annotated bounding boxes in the test data and the dotted lines are the distributions of the predicted bounding boxes at different input image scales.

**Figure S.9:**
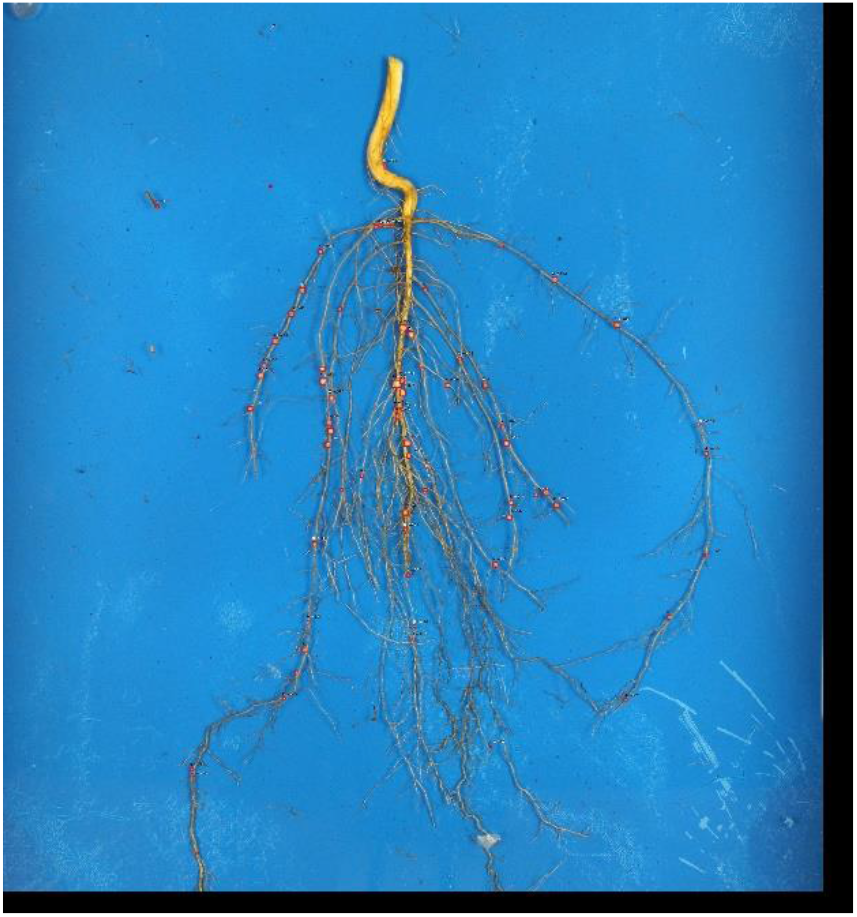
Representative example of good nodule detection on a V5 growth stage soybean root.

**Figure S.10:**
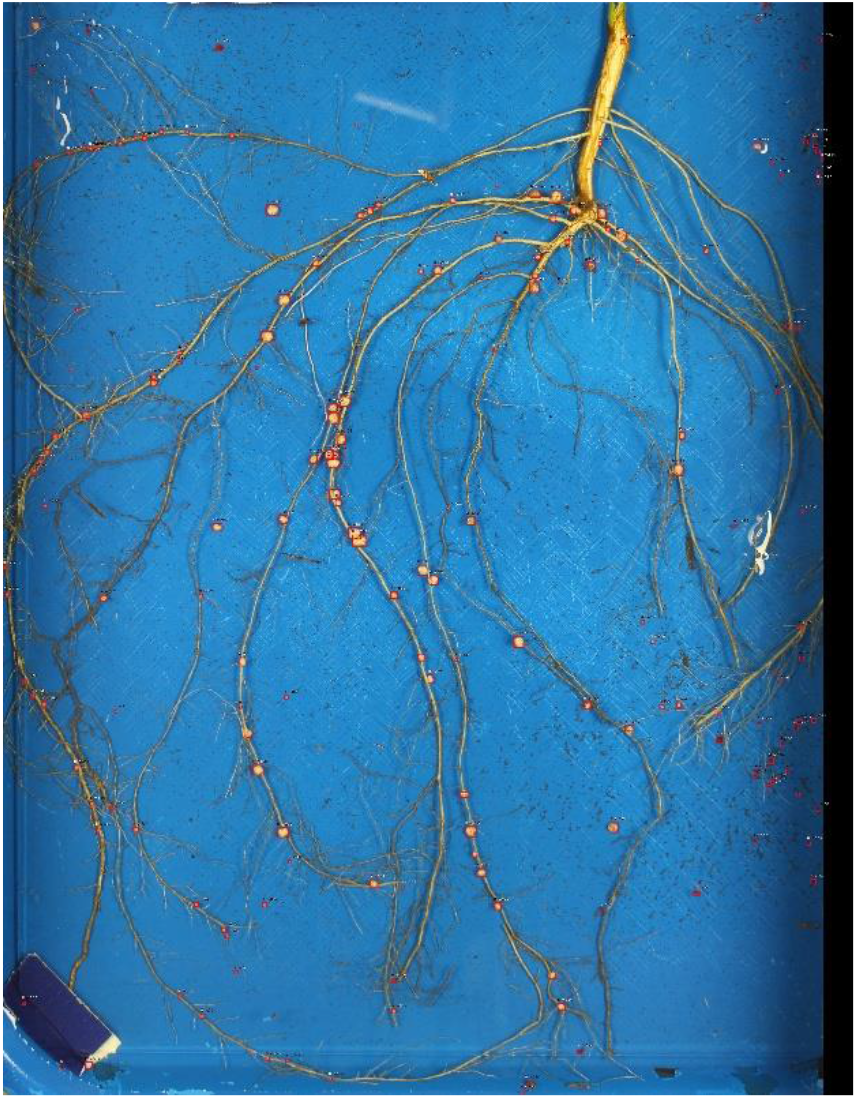
A rare example of high misclassification of image debris as nodules on a V5 growth stage soybean root. While most of the actual nodules were accurately identified, additional materials in the image were misclassified as nodules.

**Table S.1:**
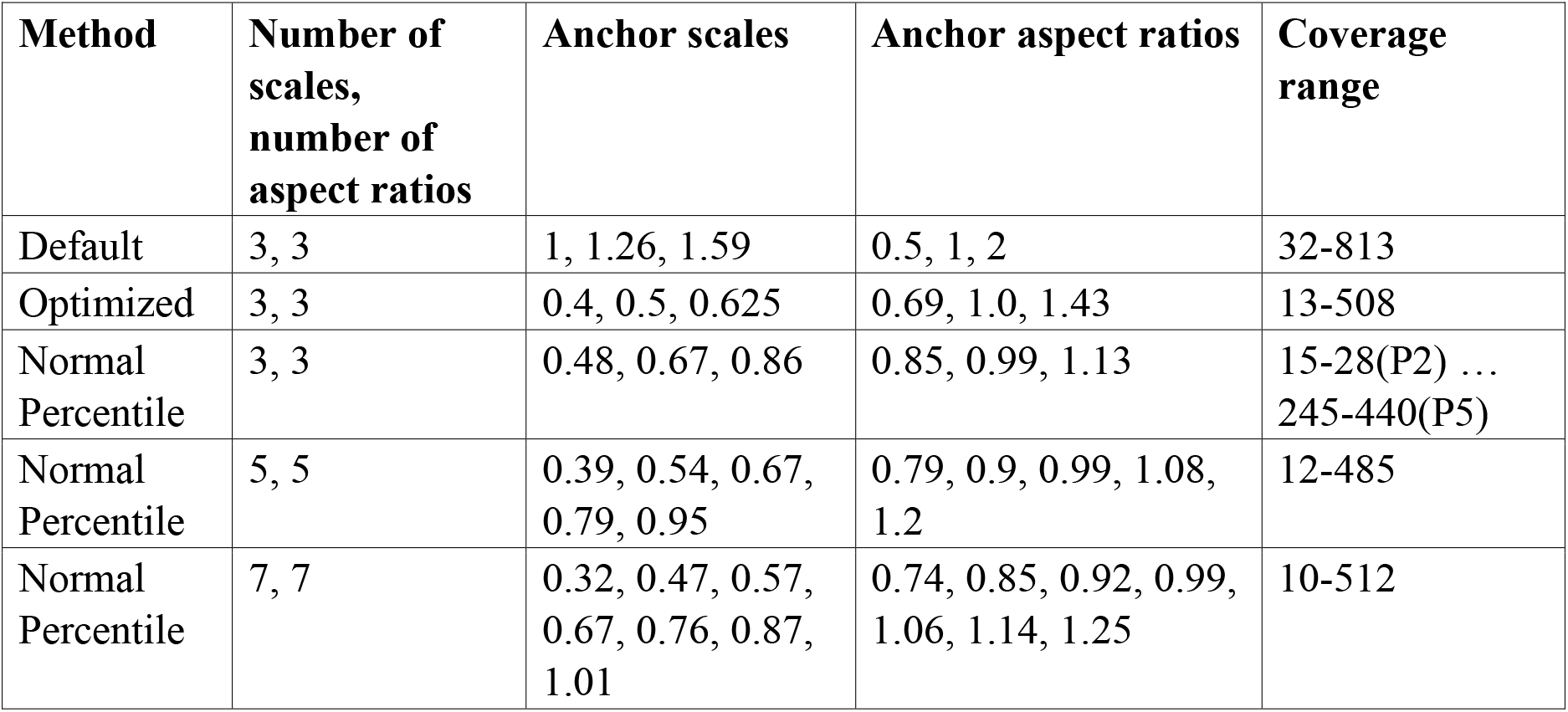
Sizes, aspect ratios, and range of anchor configuration.

**Table S.2:** Comparison of average times taken to extract, wash and image roots in this study with average times required to hand quantify nodules compared to SNAP Quantify.

| Growth Stage | Average time required in seconds for tasks: |  |  |  | Factors of SNAP Improvement Over Hand Quantification |
| --- | --- | --- | --- | --- | --- |
|  | Extraction | Washing & Image | Hand Quantify | SNAP Quantify |  |
| V1 | 240 | 100 | 1500 | 90 | 16.6 x |
| V3 | 360 | 128 | 2100 | 120 | 17.5 x |
| V5 | 420 | 150 | 3000 | 150 | 20.0 x |

